# Cohesin-mediated genome architecture does not define DNA replication timing domains

**DOI:** 10.1101/542902

**Authors:** Phoebe Oldach, Conrad A. Nieduszynski

## Abstract

3D genome organization is strongly predictive of DNA replication timing in mammalian cells. This work tested the extent to which loop-based genome architecture acts as a regulatory unit of replication timing by using an auxin-inducible system for acute cohesin ablation. Cohesin ablation in a population of cells in asynchronous culture was shown not to disrupt patterns of replication timing as assayed by replication sequencing (RepliSeq) or BrdU-focus microscopy. Furthermore, cohesin ablation prior to S phase entry in synchronized cells was similarly shown to not impact replication timing patterns. These results suggest that cohesin-mediated genome architecture is not required for the execution of replication timing patterns in S phase, nor for the establishment of replication timing domains in G1.

## 1. Introduction

The double helix structure of DNA immediately lends itself to a mechanism for genome replication so simple and so elegant that the question of faithful transmission of hereditary information, to some, seemed solved as soon as it was posed [1]. But behind the elegance of the underlying mechanism, the process of genome replication is in fact a complex, high-stakes, and intricately regulated cellular undertaking. A significant factor necessitating regulation is the sheer scale of the process: in a human cell, 12 billion bases are replicated in about 8 hours, with replication initiating from somewhere between 12,000 and 250,000 replication origins spread across 23 chromosome pairs [2]. Intriguingly, not all of these origins fire in any given S phase. Of those that do fire, not all fire at the same time. Instead, clusters of contiguous origins appear to fire in synchrony, leading to zones of hundreds of kilobases to megabases of DNA making up distinct replication timing domains (RD) [3]. The patterns of replication timing domains, driven by selective origin firings, lead to nuanced genome-wide replication timing profiles distinct to cell type and differentiation status [4–7].

Work to characterize the cellular determinants of replication timing has uncovered a strong correlation to genome organization. This has been assayed through both cytological studies of replication foci over the course of S phase [8–11] and chromosome conformation capture studies [12–15]. Genome structure appears primarily to emerge from two fundamental organizing principles:(1) a loop-based organization dependent upon the proteins cohesin and CTCF results in domains of ≤1 Mb that are defined by enriched internal interactions and known as topologically associating domains (TADs) [16–18] and (2) compartmentalization into like chromatin states results in alternating 1-10 Mb domains of active or inactive chromatin [16,19,20]. Of these different classes of genome structure, there is evidence that replication timing domains align well with compartments: early-replicating loci are significantly enriched for marks of open and active chromatin, or compartment A, whereas late-replicating loci share marks with compartment B [3,21].

Beyond the concordance between replication timing domains and genome compartments, the smaller TADs offer a compelling possibility for a unit of replication timing domain regulation. Their boundaries were found to map almost one-to-one to replication timing domain boundaries [6], and cohesin was found to be enriched at origins as well as at TAD boundaries [22]. A link between TAD-based genome structure and replication origin function was further supported by the finding that both are established concomitantly in early G1 [23,24] and change correspondingly over the course of cellular differentiation [25]. The loop-based mechanism by which cohesin is hypothesized to drive genome organization is furthermore in agreement with work showing a correlation between chromatin loop size and interorigin distance: the relationship held as reprogramming of differentiated cells to embryonic cells resulted in both a shortening of interorigin distances and chromatin loop domains [26]. Together, this abundance of correlative evidence has made a convincing case for a ‘Replication Domain Model’ in which the stable and spatially distinct structural unit of TADs gives rise to the stable functional unit of replication timing domains [6,27].

Due to cohesin’s critical role in chromosome segregation, studies of the impact of cohesin on genome architecture and replication timing have until recently been limited by the inability to carry out a clean cohesin knock-out or complete depletion over the course of a single cell cycle. A study which used cohesin knockdown by siRNA showed a reduction in active origins and an increase in interorigin distance in response to depleted cohesin levels, however this work may have been limited by the slower approach of regulating cohesin at the transcriptional level, and furthermore did not explore an effect on replication timing domains [22]. The development of the auxin-inducible degron system [28,29] has made this sort of research much more tractable, allowing for a rapid and near-complete depletion of cohesin. Chromosome conformation studies of cohesin ablation over a single cell cycle via a degron tag on cohesin’s Scc1 subunit, have shown a rapid loss of TADs, while compartment structure remained intact [30].

In this study, an auxin-inducible degron tag on the Scc1 subunit of cohesin is exploited to test whether the observation of a strong correlation between cohesin-mediated genome architecture and replication timing is substantiated as a causative relationship.

## 2. Materials and Methods

### 2.1. Cell culture

HCT116 cells [29] were grown in MyCoys 5A Media with 10% FBS, 100 U/mL penicillin, and 100 µg/mL streptomycin at 37°C with 5% CO_2_. Cells were treated with 500 µM indole-3-acetic acid (IAA, Sigma I5148) to induce loss of mAID-tagged Scc1.

### 2.2. Cell Sorting

Cell sorting was performed as described previously [31]. In brief, asynchronous cells were treated or not treated with IAA (500 µM) for 2 hours and then treated with BrdU (100 µM) for 2 hours. Cells were washed, trypsinized and harvested, and fixed with 75% ethanol. Fixed cells were prepared for fluorescence activated cell sorting (FACS) with RNase A (250 µg/mL) and propidium iodide (PI, 50 µg/mL). Cells were sorted on the basis of PI stain into 4 bins spanning S phase to collect a minimum of 2.5×10^5^ cells per bin.

### 2.3. Synchronization

Cells were treated with 2.5 mM thymidine for 24 hours and released by washing into fresh media. 3 hours after release, cells were treated with 100 ng/mL nocodazole for 8 hours, at which point mitotic cells were collected by shake-off. Cells were plated into fresh media without nocodazole and 30 minutes later were treated with 500 µM auxin or left untreated. Cells were collected at 4, 7, or 10 hours post release from nocodazole, and BrdU was added to 100 µM 2 hours before the collection of each cell population (*i.e.* at 2, 5, and 8 hours).

### 2.4. Flow cytometry

To confirm loss of Scc1 upon auxin treatment cells were fixed with 75% ethanol and prepared for flow cytometry assessment of mClover signal.

To confirm the cell cycle staging of BrdU-pulsed synchronized samples, cells were fixed and permeabilized with 75% ethanol. DNA was denatured by incubation in 0.2 mg/mL pepsin in 2 M HCl for 20 minutes. Cells were washed twice with wash solution composed of : 1% BSA/0.5% (v/v) Tween 20 in Dulbecco’s phosphate buffered saline (dPBS). Cells were then incubated in wash solution with anti-BrdU antibody (BD347580) at 1:200 for 1 hour. Cells were washed twice then incubated with AlexaFluor 647 Goat anti-Mouse (Invitrogen A32728) antibody at 1:100 in wash solution for 1 hour, followed by two washes. Cells were finally treated with RNase A (250 µg/mL) and PI (50 µg/mL) before acquisition on a BD LSRFortessa.

### 2.5. RepliSeq

DNA was extracted from each sample using phenol/chloroform followed by an isopropanol precipitation. The extracted DNA was sonicated to an average fragment size of 300 bases. DNA was purified using SPRI beads, repaired with the NEBNext End Prep kit followed by ligation of sequencing adaptors using the NEBNext UltraII ligation kit. DNA was purified using AMPure XP beads (Beckman Coulter). BrdU pulldown was carried out as per Peace *et al*, 2016 [32]. Briefly, DNA was heat-denatured and snap cooled, then incubated overnight with anti-BrdU antibody (BD347580). The sample was then incubated for 1 hour with Protein G Dynabeads (ThermoFisher), washed three times with IP buffer (PBS/0.0625% v/v Triton X-100) and once with TE, and then eluted into TE/1% (w/v) SDS at 65°C. Eluted DNA was cleaned using AMPure XP beads, and then amplified with Illumina indexing primers using 16 cycles of the NEBBext Ultra II kit. Amplified libraries were cleaned, quantified using NEBNext Library Quant kit, and checked for fragment length using Tapestation. Libraries were sequenced using 75 cycles on an Illumina NextSeq 500 for a minimum of ten million reads per library.

Sequenced HCT116 raw fastq files and bigwig files reporting processed log_2_ replication timing scores are available from the NCBI GEO database (accession number GSE124025). The processed replication timing scores can also be visualized via a UCSC genome browser hub : https://ln1.path.ox.ac.uk/groups/nieduszynski/Oldach2019/Oldach2019_hub.txt.

### 2.6. Replication Foci IF

#### Asynchronous

Cells were seeded onto coverslips (thickness #1.5) and grown to 80% confluency. Media was replaced and cells were treated with IAA (500 µM) or left untreated for 3.5 hours, at the end of which all cells were pulsed with BrdU (50 µM) for 30 minutes.

Upon completion of the BrdU pulse, coverslips were washed with PBS and fixed with 3% (w/v) formaldehyde/dPBS, washed, and then permeabilized with 0.2% (v/v) Triton X-100 in dPBS. DNA was denatured with 4 M HCl treatment for 10 minutes, after which coverslips were washed and blocked with 5% (w/v) BSA in PBST. Cells were incubated for 1 hour with anti-BrdU antibody (BD347580) 1:200 in block solution. After three washes, coverslips were incubated for 1 hour in the dark with AlexaFluor Rabbit anti Mouse 568 (Invitrogen A11061) at 1:1000 in 5% BSA. Coverslips were washed, and nuclei were stained with DAPI. Coverslips were finally mounted with VectaShield H1000. DAPI signal was used to identify cells for image acquisition, and BrdU-positive cells were classified by eye into five S phase stages by researchers blinded to the treatment status of the images [9].

#### Scc1 IF

Asynchronous cultures of HCT116-Scc1-mAID cells were seeded on coverslips, and treated for 2 hours with IAA (500 µM) or left as controls. At the end of the treatment, coverslips were washed with PBS then fixed with 3% formaldehyde. Cells were permeabilized with 0.2% Triton X-100 in PBS, washed with PBST, then blocked for 1 hour in 5% BSA in PBST. Coverslips were incubated for 1 hour in anti-Scc1(Abcam 154769) at 1:500 in block. After three washes coverslips were incubated for 1 hour in Alexafluor 647 Goat anti Rabbit (Invitrogen A21245) 1:1000 in block. Coverslips were washed, stained with DAPI, and mounted in Vectashield.

### 2.7. RepliSeq Computational Analysis

Reads were aligned to the human genome (hg38) with STAR [33]. Uniquely mapping reads were binned into 1 kb windows and these windows were filtered to remove bins with signal spikes (Z score > 2.5). Reads were then binned into 50 kb windows. E/L RT was calculated as the log_2_(S phase bin 1/S phase bin 3) reads for the asynchronous sample and log_2_(4h/10h) for synchronized replicates. Quantile normalization and Loess smoothing with a span of 300 kb were applied to E/L RT values. RepliSeq data for wildtype HCT116 cells (Accessions: 4DNFI5BZJXDE, 4DNFIC4VUF86) and IMR90 cells (Accessions: 4DNFILOYZWEM, 4DNFIKQDQCNB) were gathered from the 4D Nucleome data portal [34], and a Pearson correlation was used to quantify the degree of similarity between the profiles.

A Z score was calculated based on the difference between treated and control replication timings to identify significantly changed bins. An FDR adjustment was applied to *p* values to correct for multiple testing.

Weighted RT values were calculated as the weighted average of the four S phase bins for asynchronous samples, or three S phase bins for synchronized samples. Publicly accessible data for histone marks, CTCF binding sites, and other genome characteristics in HCT116 cells was extracted from ENCODE [35] and the UCSC Table Browser [36] (Supplemental Table 1). ProSeq data from Rao *et al*, 2017 was used to identify genes with differential expression in response to cohesin loss upon auxin treatment. Briefly, significantly changed genes were classified as those with an absolute fold change greater than two, an adjusted *p* value less than 0.05, and at least 0.5 RPKM. These cutoffs yielded 74 genes which were induced upon auxin treatment and 16 which were downregulated. The genomic coordinates for these genes was extracted using BioMart [38]. BEDOPS was used to calculate the mean weighted RT for auxin-treated or control samples across annotated genomic ranges [37].

**Table 1.** Genomic loci showing significant difference in both asynchronous and synchronized experiments.

| Chr | Start | End | RT Control - RT IAA |  |
| --- | --- | --- | --- | --- |
|  |  |  | Asynchronous | Synchronous |
| chr3 | 198100000 | 198150000 | -3.440337 | 3.242949 |
| chr4 | 49300000 | 49350000 | 2.723062 | 2.296154 |
| chr4 | 119400000 | 119450000 | 3.486406 | -1.410933 |
| chr4 | 190000000 | 190050000 | 4.479181 | -1.897292 |
| chr4 | 190050000 | 190100000 | 12.678844 | -5.136778 |
| chr4 | 190100000 | 190150000 | 25.976415 | -10.545368 |
| chr5 | 49650000 | 49700000 | -3.258203 | 4.272367 |
| chr5 | 181250000 | 181300000 | 2.921178 | -1.623559 |
| chr7 | 56850000 | 56900000 | 2.779265 | 1.355865 |
| chr9 | 150000 | 200000 | 3.343623 | -1.447844 |
| chr13 | 114300000 | 114350000 | 3.460791 | 1.561016 |
| chr13 | 114350000 | 114364328 | 7.856382 | 3.064328 |
| chr16 | 90100000 | 90150000 | 4.566427 | -3.221925 |
| chr21 | 6350000 | 6400000 | 3.521525 | -2.910266 |

A mappability score from Karimzadeh, *et al* [39] was used to mask unmappable regions of the genome. Analysis was carried out in R [40] and data were visualized using ggplot2 [41] and the UCSC Genome Browser [42].

The 3D genome browser [43] was used to visualize and extract TAD calls from control auxin-treated HCT116-Scc1-mAID cell HiC data from Rao *et al*, 2017, yielding a list of 2,307 domains in untreated cells and 883 domains in auxin-treated cells. Juicer [44] was used to calculate compartment boundaries from untreated HCT116-Scc1-mAID cell HiC data. TAD boundaries that fell within 500 kb of a compartment boundary were considered separately, to control for the effect of compartment status on replication timing. Compartment-independent TADs were classed as “Scc1-dependent” if they were not present (within 500 kb) in the contact domains of auxin-treated HCT116-Scc1-mAID HiC data, or “Scc1-invariant” if they remained in the contact domains of auxin-treated cells.

## 3. Results

### 3.1. Efficient and rapid degradation of endogenous Scc1

It has been previously shown that degradation of the Scc1 subunit of the cohesin complex leads to an inability of the complex to associate with DNA [30]. Here, we used an HCT116 cell line with the endogenous Scc1 tagged with mini-AID and mClover for rapid, auxin-inducible degradation of the cohesin subunit and thus destabilization of the cohesin complex [29]. To confirm loss of the Scc1 fusion protein after two hours of auxin treatment, we analyzed mClover signal in treated and control cells by microscopy (Figure 1A) and flow cytometry (Figure 1B). These two methods yielded comparable values: after two hours of auxin treatment 73% of cells were mClover-negative by flow cytometry (n=40,649), and 78% were mClover-negative by microscopy (n=54) (Figure 1B,C). Cells that retained residual mClover signal after auxin treatment had notably reduced intensity of signal relative to untreated levels (Figure 1A). To further confirm that loss of mClover related to loss of Scc1, immunofluorescence (IF) was carried out for Scc1. 81% of cells imaged were negative for Scc1 stain after two hours of auxin treatment and those cells that retained any Scc1 signal after treatment were markedly reduced in signal intensity (Figure 1A,C). These results agree with previous work that has demonstrated a loss of Scc1 within an hour of auxin treatment in the same cell line [29,30].

**Figure 1.**
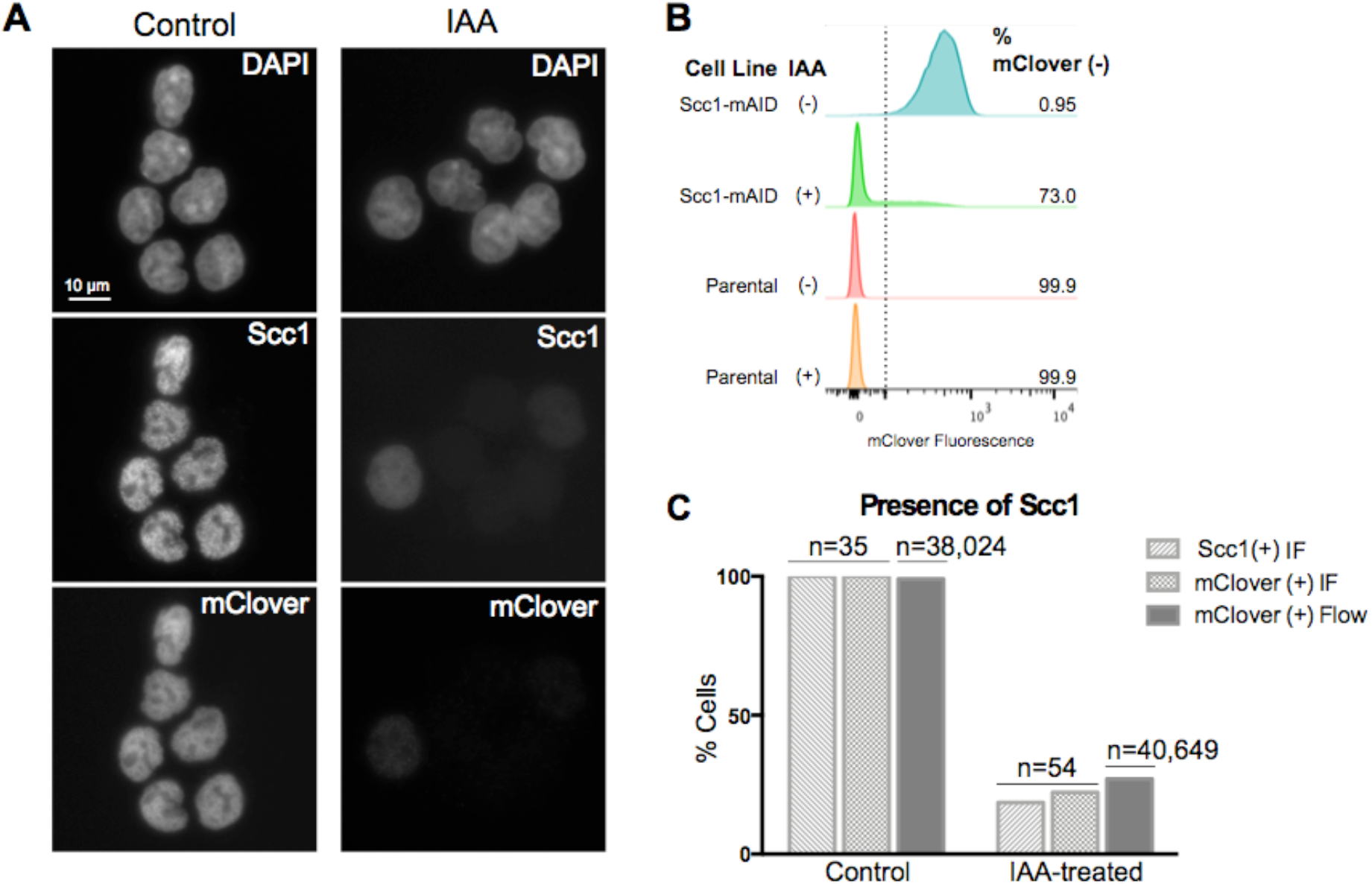
Auxin treatment results in loss of Scc1. A. Immunofluorescence for Scc1 and mClover in HCT116 Scc1-mAID-mClover cells under control conditions or two hours of auxin treatment. B. Flow cytometry analysis for mClover fluorescence using both HCT116 Scc1-mAID-mClover cell line and parental HCT116 cell line as a negative control. The mClover-positive gate was set based on the parental cell line at 10^2^ fluorescence units. C. Comparison of direct IF read-out of loss of Scc1 and indirect read out of loss of mClover through both flow cytometry and IF.

### 3.2. Acute loss of cohesin in S phase does not perturb replication patterns

To analyze the impact of cohesin loss on DNA replication timing, we have used RepliSeq on asynchronously-growing auxin-treated and untreated control cells [31,32]. Asynchronously growing cells were treated with auxin for two hours or left as untreated controls, pulsed with BrdU for a further two hours, and then FACS enriched on DNA content into four S phase fractions. DNA was extracted from each fraction, then BrdU-containing nascent DNA was immunoprecipitated and subjected to high-throughput sequencing. The proportion of nascent DNA in each fraction served as a measure of the relative replication timing across the genome. In the mock-treated samples, replication timing was found to be in good agreement with previously reported HCT116 cell replication timing data (Supplemental Figure 1) [45]. The overall profile of replication timing did not change in response to auxin-induced Scc1-ablation (Figure 2A). Quantitatively, the correlation between the control and treated profiles was comparable to the correlation between the control and a previously-reported HCT116 replication timing profile (Figure 2C). To facilitate identification of regions with changed replication timing, the difference of replication timing in control versus auxin-treated samples was calculated for each genomic locus. Replication timing was only significantly changed (FDR-adjusted p < 0.05) in 0.34% of mappable genomic bins (189 bins out of 56,034 bins with mappability score > 0.5) (Figure 2B) [39]. Significant differences were enriched in bins with low mappability and dispersed across the genome (median distance between significant bins: 925,000 bases), suggesting that these differences were a consequence of experimental noise rather than domains of biological relevance. Furthermore, loci with significantly-changed replication timing were not enriched at TAD boundaries, as might be expected if these changes resulted from loss of TAD architecture (Figure 2D).

**Figure 2.**
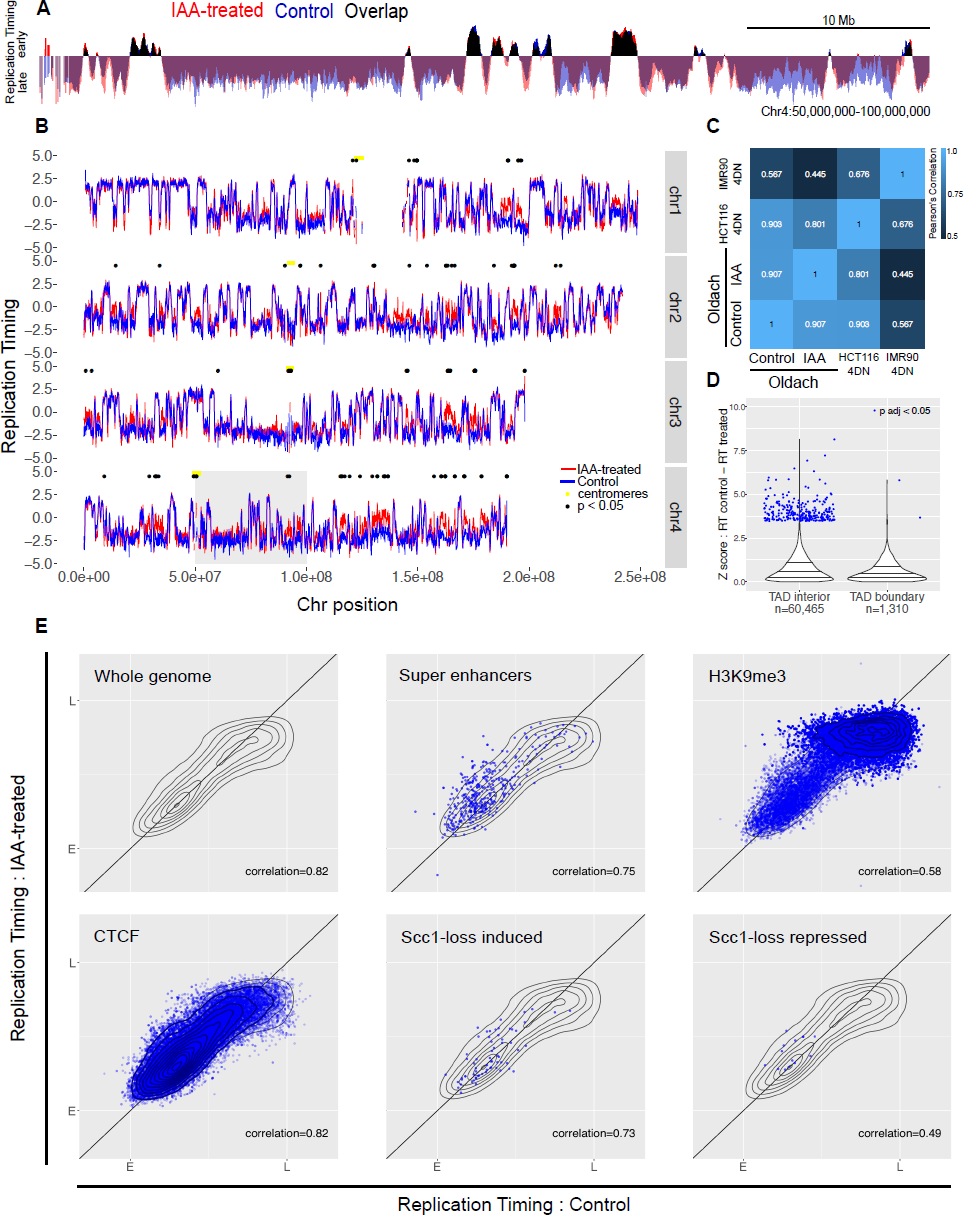
Acute loss of cohesin in S phase does not perturb replication timing. A. Example locus overlay of replication timing (RT, log-scaled ratio of read counts in early over late S phase bin) for auxin-treated versus control cells. B. RT plots for four example chromosomes, with significantly-changed loci marked with black dots. The gray box highlights the region shown in (A). C. Pearson correlation calculated for RT values across genome, compared to previously-published 4D Nucleome RepliSeq data. D. Violin plot showing the distribution of Z scores for control versus auxin-treated RT difference across the genome, separating out genomic bins covering a TAD boundary. E. RT for control versus auxin-treated cells across the whole genome, and across genomic coordinates annotated for super enhancers, H3K9me3, CTCF, and genes that showed induction or repression upon auxin treatment in the Scc1-mAID cell line. The gray topology plot in each subfigure shows the whole-genome behavior, and genome features with dense annotations (CTCF and H3K9me3) have a topology map in dark blue.

As genome-wide analysis did not demonstrate widespread perturbations to the replication timing profile in response to the loss of cohesin, genomic regions marked by features hypothesized to show a change were specifically assessed. These included genomic regions with features characteristic of early replication (super enhancers, DNase hotspots, CpG islands, H3K4 methylated regions, H3K9 and H3K27 acetylated regions), late replication (H3K9me3, polycomb binding sites), CTCF binding sites, or genes with expression changes in response to auxin-induced loss of Scc1. None of these sets of sites showed a differential response (Figure 2E, Supplemental Figure 6).

To investigate the impact of cohesin ablation on the cytological patterns of replication timing, auxin-treated or control BrdU-pulsed Scc1-mAID-mClover cells were classed into five S phase categories based on their BrdU foci patterning (Figure 3A) [9]. The treatment with auxin and resultant loss of cohesin did not alter the abundance of cells across each of the five classes (Figure 3B). In combination with the finding that auxin-induced loss of Scc1 does not change progression through S phase (Supplemental Figure 3), this suggests that auxin treatment does not change the patterns of replication characteristic to different stages of S phase.

**Figure 3.**
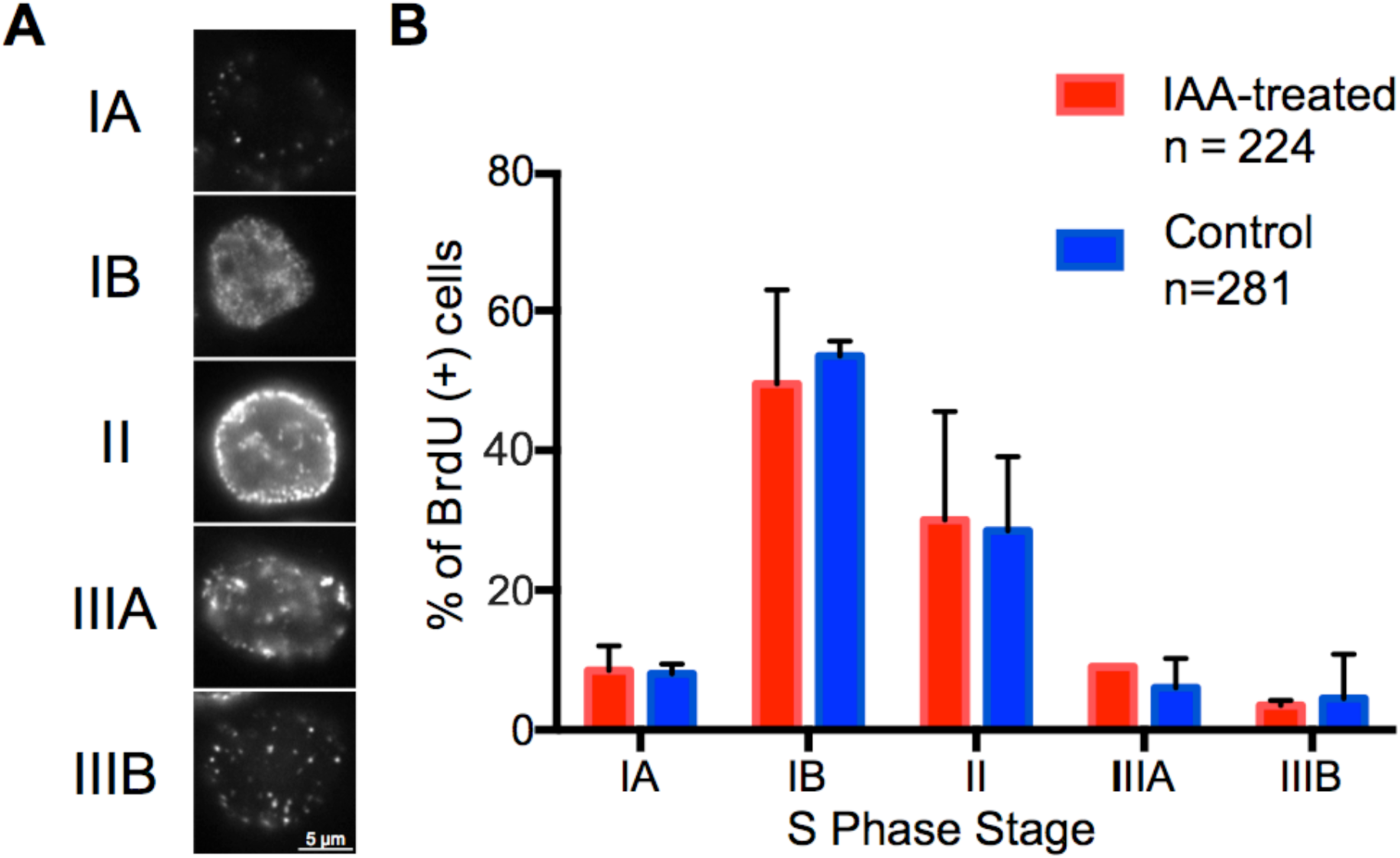
Acute loss of cohesin in S phase does not perturb patterns of replication foci. A. Replication foci patterns were assayed using BrdU IF in asynchronous cells after auxin or control treatment. B. Number of cells in each S phase stage were counted for two biological repeats of auxin-treated and untreated samples.

### 3.3. Loss of cohesin from early G1 does not perturb replication timing

To assess whether cohesin is required for the establishment, rather than execution, of replication timing, it was necessary to ablate cohesin from cells before the onset of S phase. This also diminished the potential for artifacts arising from the impact of cohesin ablation on chromosome segregation. Cells were synchronized with a 24 hour thymidine block and released for three hours, followed by nocodazole treatment for eight hours. Cells were released from nocodazole by mitotic shake-off and 30 minutes after the shake-off auxin was introduced for treated cells. Both auxin-treated and untreated cells showed similar cell cycle progression following release (Supplemental Figure 3).

Synchronized auxin-treated or untreated control cells were pulsed with BrdU at two, five, and eight hours post-release. Two hours after the addition of BrdU to each synchronized timepoint the cells were harvested and DNA was extracted. BrdU-labeled nascent DNA was immunoprecipitated and subjected to high-throughput sequencing. The overall replication timing profiles for synchronized cells with cohesin depleted from early G1 showed the same pattern to those of untreated synchronized cells (Figure 4A). The variability between treated and control cells was comparable to that between control cells and an independent control HCT116 dataset (Figure 4C). 219 bins showed significant changes in response to cohesin ablation (out of 56,034 50 kb bins with map score > 0.5), and these were spread across the genome (median distance between bins 700,000 bases) and enriched in unmappable regions such as centromeres (Figure 4B). Just 14 of these bins were shared with the significant bins from the asynchronous sample, and in four of these bins the direction of the change was different – suggestive of noisy or unmappable loci rather than a consistent biological effect (Table 1).

**Figure 4.**
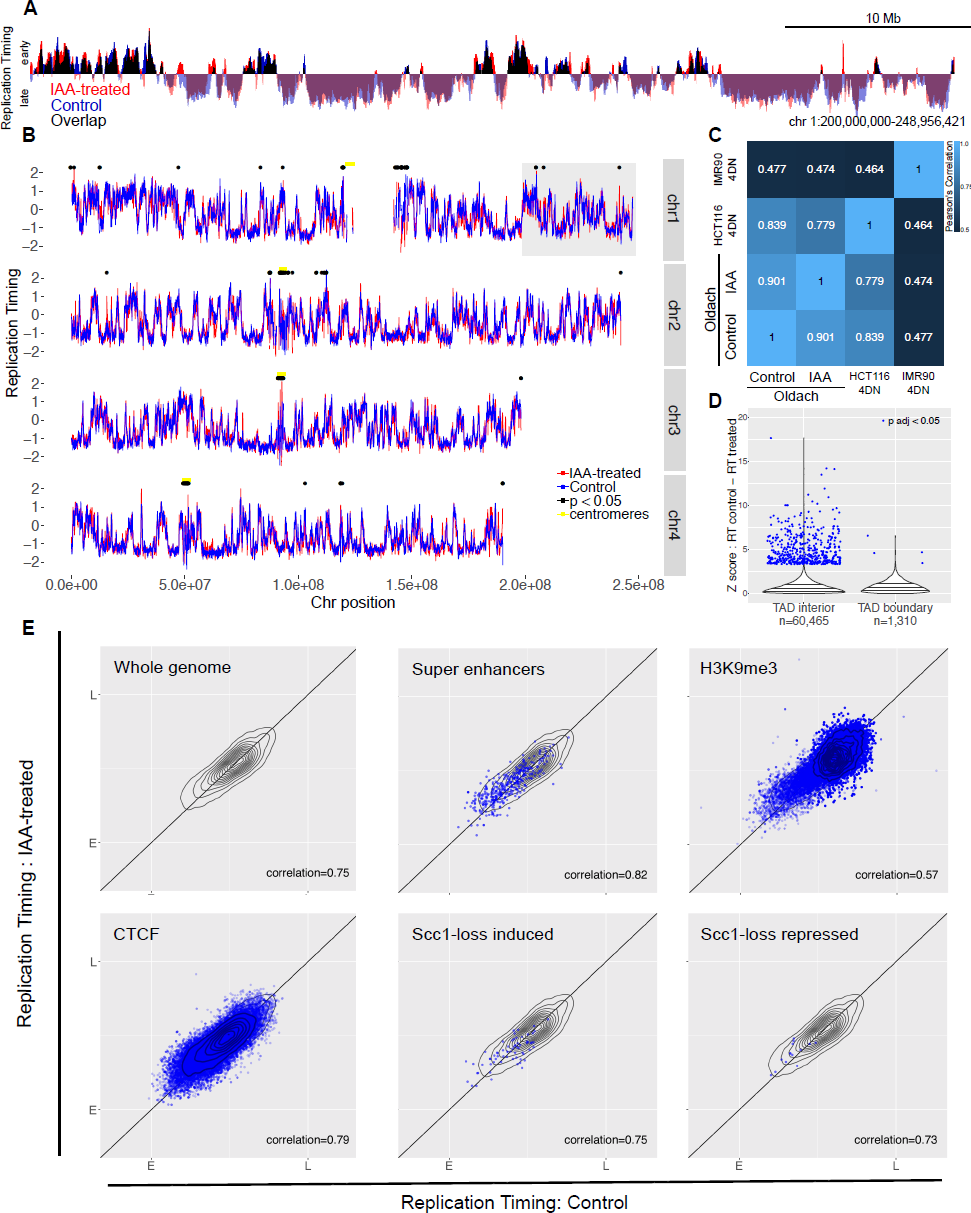
Loss of cohesin from early G1 does not perturb replication timing. A. Example locus overlay of replication timing (RT, log-scaled ratio of four hour timepoint read counts over ten hour timepoint counts) for auxin-treated versus control cells. B. RT plots for four example chromosomes, with significantly-changed loci marked with black dots. The gray box highlights the region shown in (A). C. Pearson correlation calculated for RT values across genome, compared to previously-published 4D Nucleome RepliSeq data. D. Violin plot showing the distribution of Z scores for control versus auxin-treated RT difference across the genome, separating out genomic bins covering a TAD boundary. E. RT for control versus auxin-treated cells across the whole genome, and across genomic coordinates annotated for super enhancers, H3K9me3, CTCF, and genes that showed induction or repression upon auxin treatment in the Scc1-mAID cell line. The gray topology plot in each subfigure shows the whole-genome behavior, and genome features with dense annotations (CTCF and H3K9me3) have a topology map in dark blue.

As with the replication timing data from asynchronous cells, no specific genomic features showed a particular change in replication timing (Figure 4E).

### 3.4. Impact of genome organization on replication timing

The lack of response of replication timing domains to cohesin ablation led to the hypothesis that compartment boundaries may be a more robust delineator of replication timing than TAD boundaries. To assess this possibility, TAD calls from the control and auxin-treated cells of Rao *et al*, 2017 were collected from the 3D Genome Browser [43]. Upon cohesin ablation, Rao *et al* saw a pervasive loss of TAD structure while higher level compartment structure remained (Figure 5A) [30]. In keeping with this, of 2,026 compartment-independent TAD boundaries (*i.e.* not within 500 kb of a compartment boundary) mapped in the untreated HCT116 cells, only 887 remained following auxin treatment. TAD boundaries that disappeared in the auxin-treated dataset were classed as ‘Scc1-dependent boundaries’ (n=1,139), while those that were shared (within 500 kb) between control and auxin-treated datasets were classed as ‘invariant’ (n=887), and any that fell within 500 kb of an A/B compartment domain boundary were analyzed separately (n=1,378).

**Figure 5.**
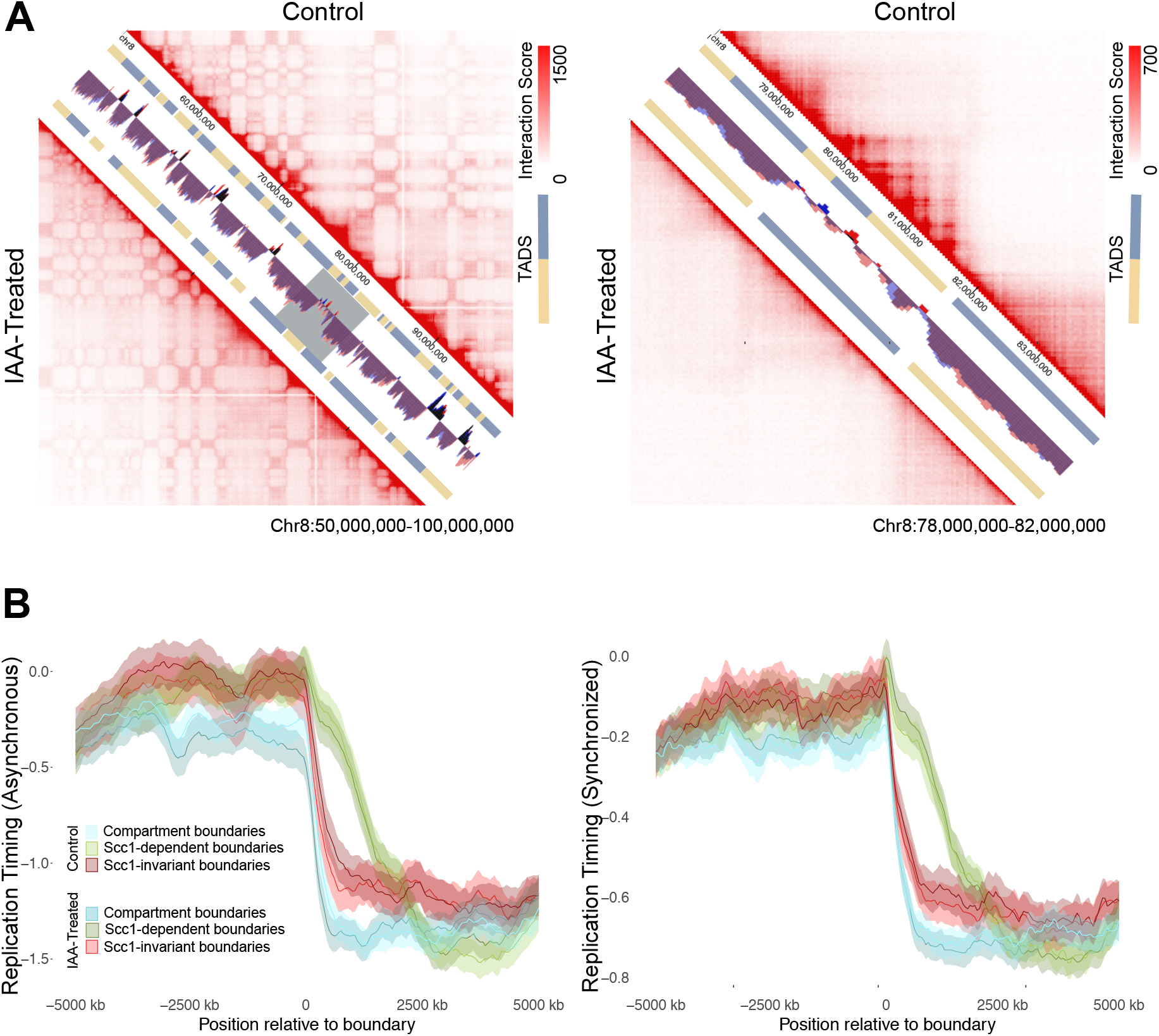
Replication timing across TAD boundaries versus compartment boundaries. A. Visualization of HiC data from Rao *et al*, 2017 in untreated cells versus auxin-treated cells. The left panel spans 50 Mb to highlight compartments and the right zooms in to 6 Mb from within that locus (gray box) to highlight changes to TAD structure. B. Replication timing metaplots for windows 5 Mb on either side of TAD boundaries. Invariant boundaries are those called in both control and auxin-treated cells, Scc1-dependent boundaries are those which were lost upon auxin treatment.

Replication timing values for auxin-treated and control cells were assessed along 10 Mb windows to either side of Scc1-invariant and dependent boundaries. Both control and auxin-treated datasets showed a similar relationship between replication timing domains and TAD boundaries: the loss of TAD organization did not lead to changes in replication timing across TAD boundaries (Figure 5B). Furthermore, Scc1-invariant and compartment boundaries were found to show a stronger relationship to replication timing domains than Scc1-dependent boundaries, as evidenced by a sharper change in replication timing at the boundary (Figure 5B).

## 4. Discussion

The hypothesis that TADs act as a fundamental unit of organization for replication timing has gained widespread support due to the near one-to-one mapping of replication timing domains and TADs [6], the concurrent establishment of both domains in early G1 [24], and their coordinated changes over the course of cellular differentiation [13,25]. Using the Scc1-degron system, we were able to explicitly test whether the loss of cohesin resulted in a perturbation to replication timing patterns. We found that neither acute loss of cohesin in S phase nor prolonged loss of cohesin from early G1 resulted in significant changes to replication timing patterns. Thus, cohesin is not required for the execution nor establishment of replication timing domains. This is in good agreement with recent work showing that depletion of TAD boundary protein CTCF also does not result in significant genome-wide perturbations to replication timing patterns [46].

The RepliSeq technique specifically informs about replication timing domains and does not shed light on specific origin usage. Thus, it remains feasible that cohesin-dependent structure does in fact play a role in the specific choice of sites of origin firing within a domain, while the timing of higher-order domains remains independent of TAD structure. This could reconcile the apparent lack of domain timing change to the changes in interorigin distance seen previously in cohesin knockdown experimentation [22].

The lack of response of replication timing domains to cohesin loss and the stronger relationship seen between cohesin-invariant and compartment boundaries than cohesin-dependent boundaries to changes in replication timing suggests that replication timing domains may be regulated by the same organizing principles that drive A/B compartmentalization rather than TADs. While this could in part be explained by the fact that neighboring TADs could share similar chromatin states – and thus may be likely to replicate at similar times regardless of being members of distinct regulatory modules – there is increasing evidence to support a model in which compartmentalization-based structure rather than loop-based structure drives replication timing. Recently-discovered early replication control elements (ERCE) have been shown to drive robust CTCF-independent interactions in a manner akin to compartmentalization, and their deletion leads to changes in replication timing [46]. Further work characterizing the cellular mechanisms driving genome compartmentalization will open pathways to perturb this system and elucidate the causal relationship between replication timing and genome topology.

## Author Contributions

conceptualization, P.O. and C.A.N.; investigation, P.O.; writing—original draft preparation, P.O.; writing—review and editing, C.A.N..; supervision, C.A.N.

## Funding

This work was supported by Biotechnology and Biological Sciences Research Council grants BB/N016858/1 and Wellcome Trust Investigator Award 110064/Z/15/Z. PFO is generously supported by a Wellcome Trust studentship award 203727/Z/16/Z.

## Acknowledgments

The authors are very grateful to Masato Kanemaki for sharing the Scc1-AC Tir1 HCT116 cell line. We thank David Gilbert for sharing unpublished results. We thank Line Eriksen and Michal Maj of the Dunn School Flow Cytometry Suite for facilitating FACS. We thank Amanda Williams and Rebecca Busby for their support at the Zoology Sequencing Facility. We would in particular like to thank the Nieduszynski lab for helpful discussions and substantial support.

## Conflicts of Interest

The authors declare no conflict of interest.

