## Supplemental Figures for "Cohesin-mediated genome architecture does not define DNA replication timing domains"

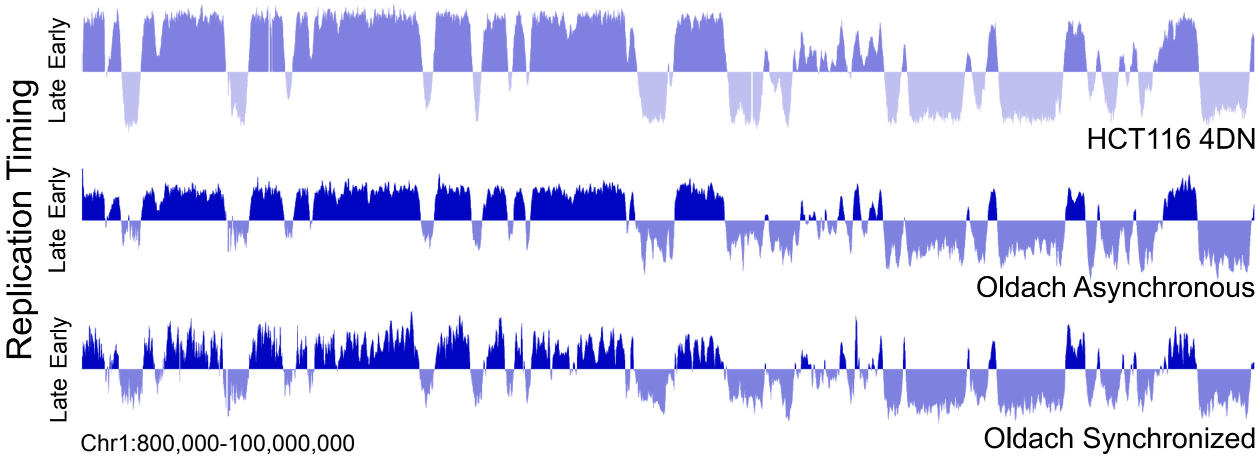


**Supplemental Figure 1. Newly generated RepliSeq data agrees with previously published RepliSeq data for HCT116 cells.** Example locus overlay of replication timing (RT, log-scaled ratio of read counts in early over late S phase bins) for untreated HCT116 cells from the asynchronous and synchronized experiments, compared to independent HCT116 data from the 4D Nucleome portal.

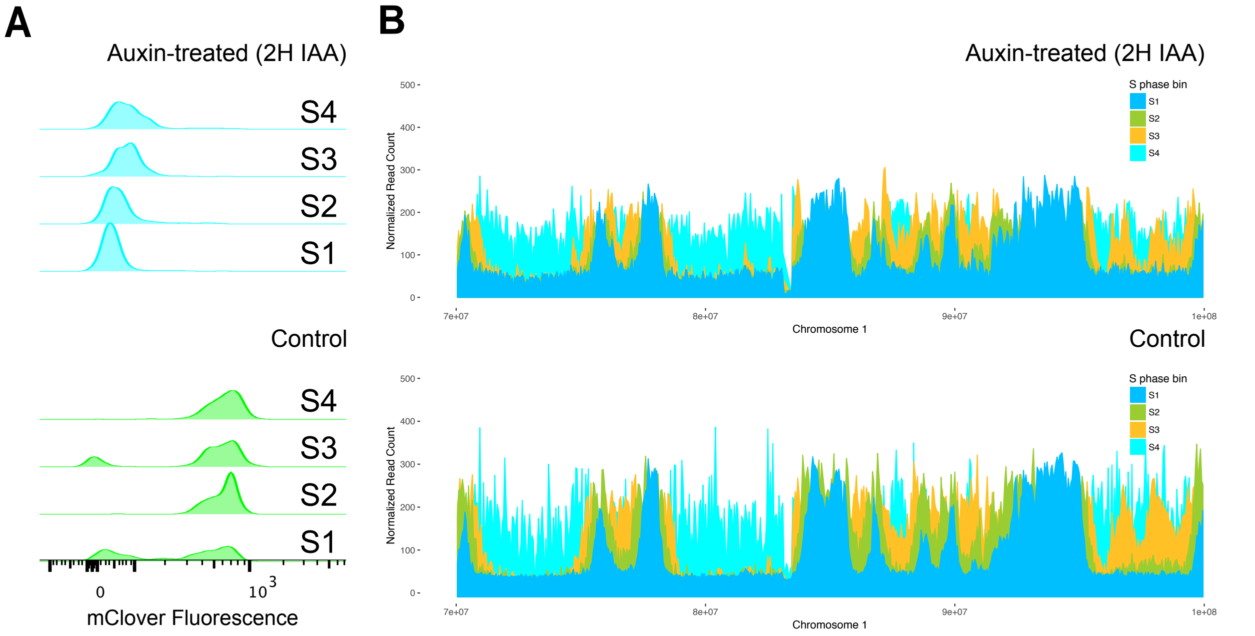


**Supplemental Figure 2. Sample preparation for asynchronous RepliSeq experiment.** A. Auxin-treated cells showed loss of cohesin as assayed by loss of mClover signal. B. In asynchronous sorted samples BrdU incorporation proceeds outward from specific peaks in the early S phase (S1) sample to pan-genome incorporation in the late S (S4) sample.

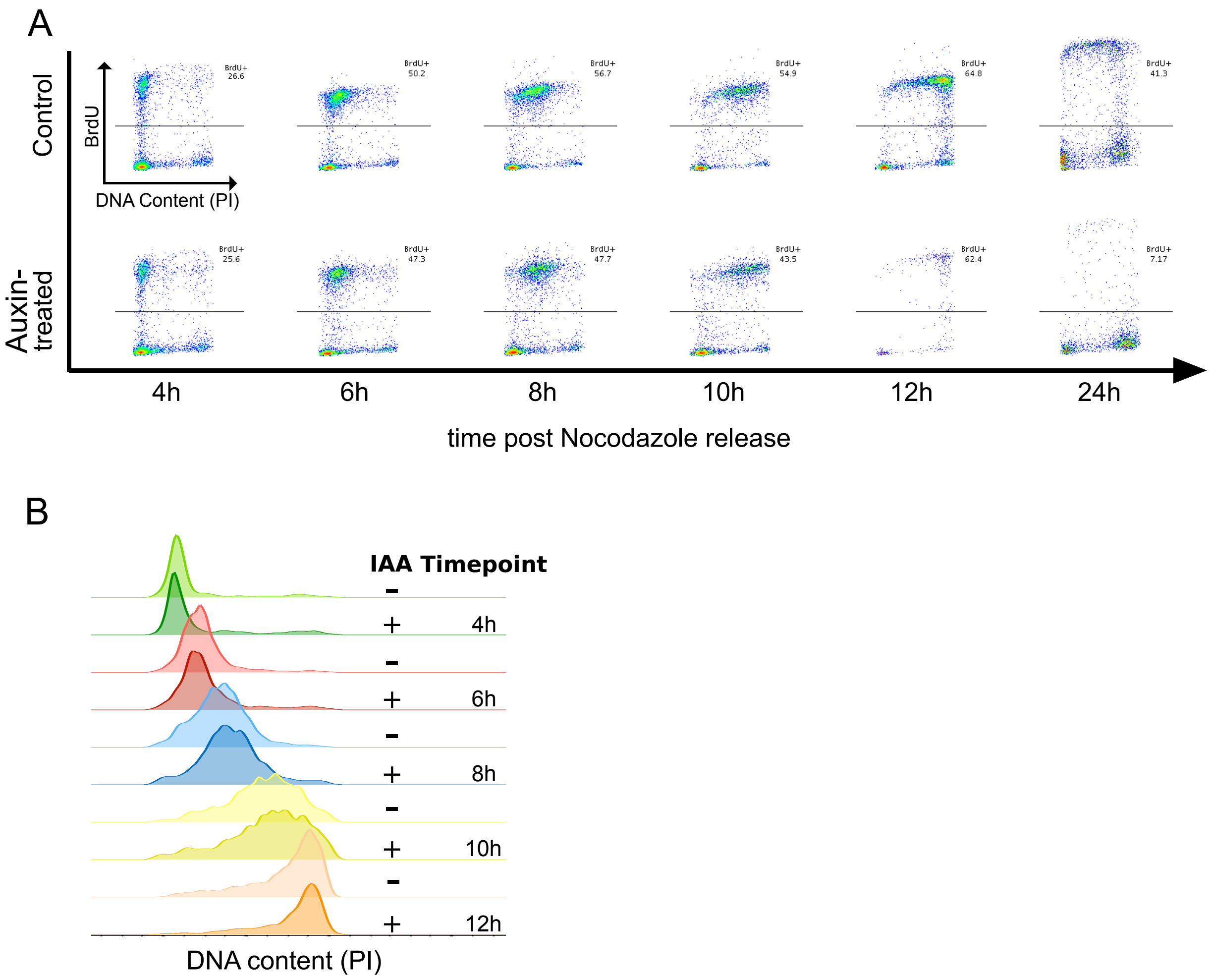


**Supplemental Figure 3. Auxin treatment post release from nocodazole by mitotic shake-off does not perturb synchronous progression through S phase.** A. Flow cytometry data for DNA content versus BrdU signal showing progression through S phase upon release from nocodazole by mitotic shake-off. B. DNA content as quantified by PI signal for BrdU (+) gated cell populations of auxin-treated or control samples at varying timepoints following mitotic shake-off.

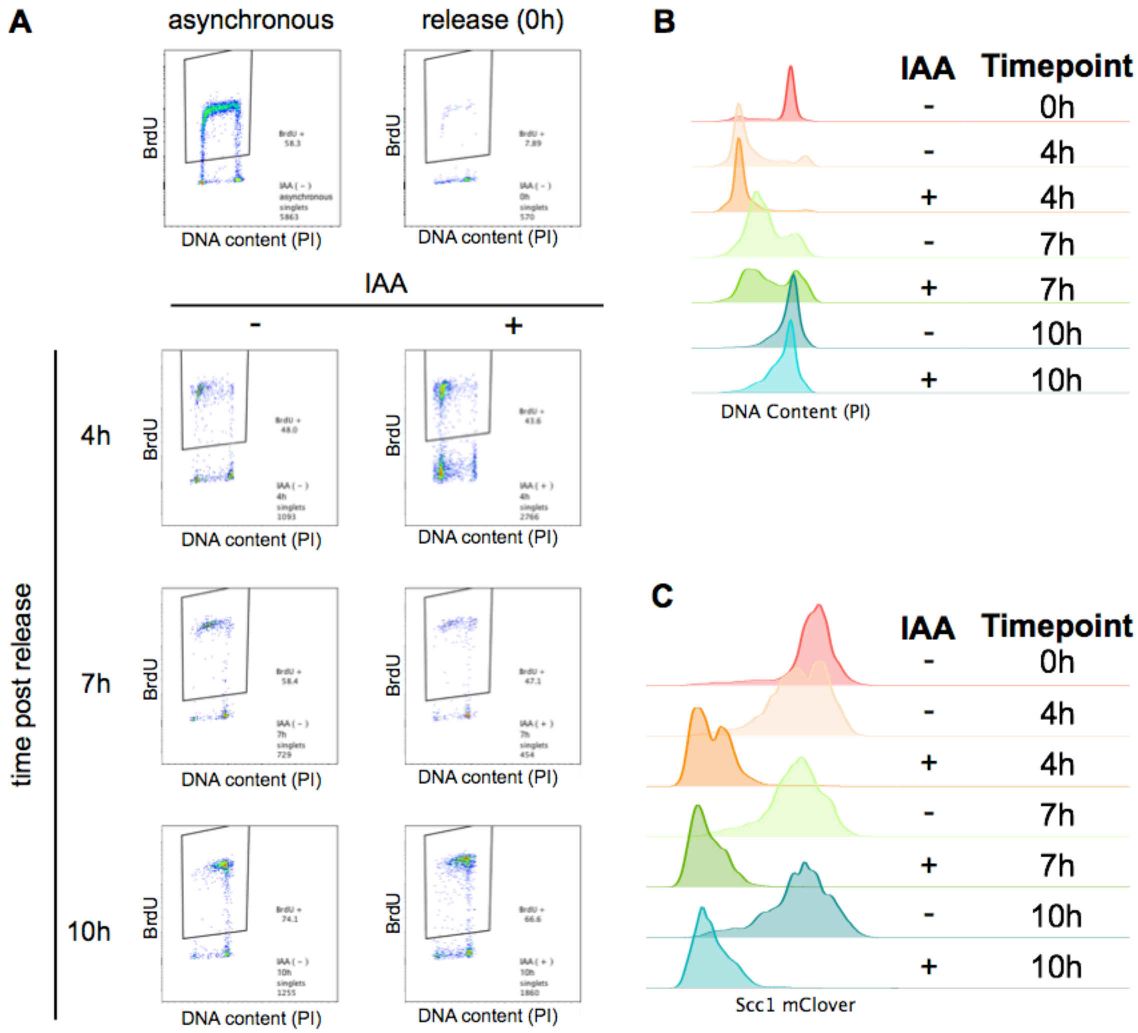


**Supplemental Figure 4. Sample preparation for synchronous RepliSeq experiment** A. Samples were collected at 4, 7, and 10 hours post-release to assess replication timing in early, mid, and late S phase, respectively. B. Flow profile gated on BrdU (+) singlets show synchronous progression through S phase. (0h post release sample not gated for BrdU (+) cells). C. Loss of Scc1 in synchronized samples was confirmed via loss of mClover signal.

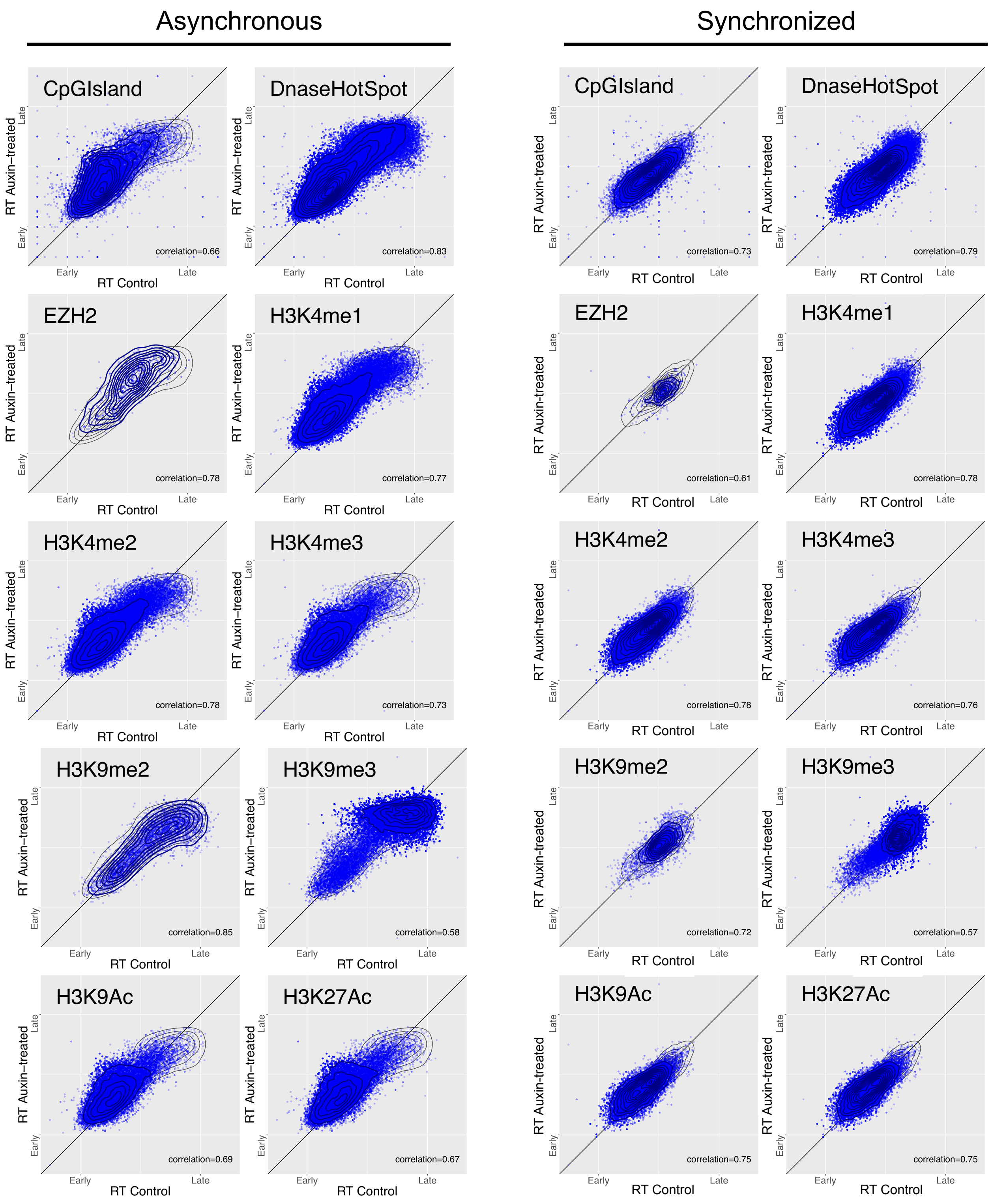


**Supplemental Figure 5.** Additional plots of control versus auxin-treated replication timing at genomic loci with chromatin states or protein binding sites associated with characteristically early or late replication timing.

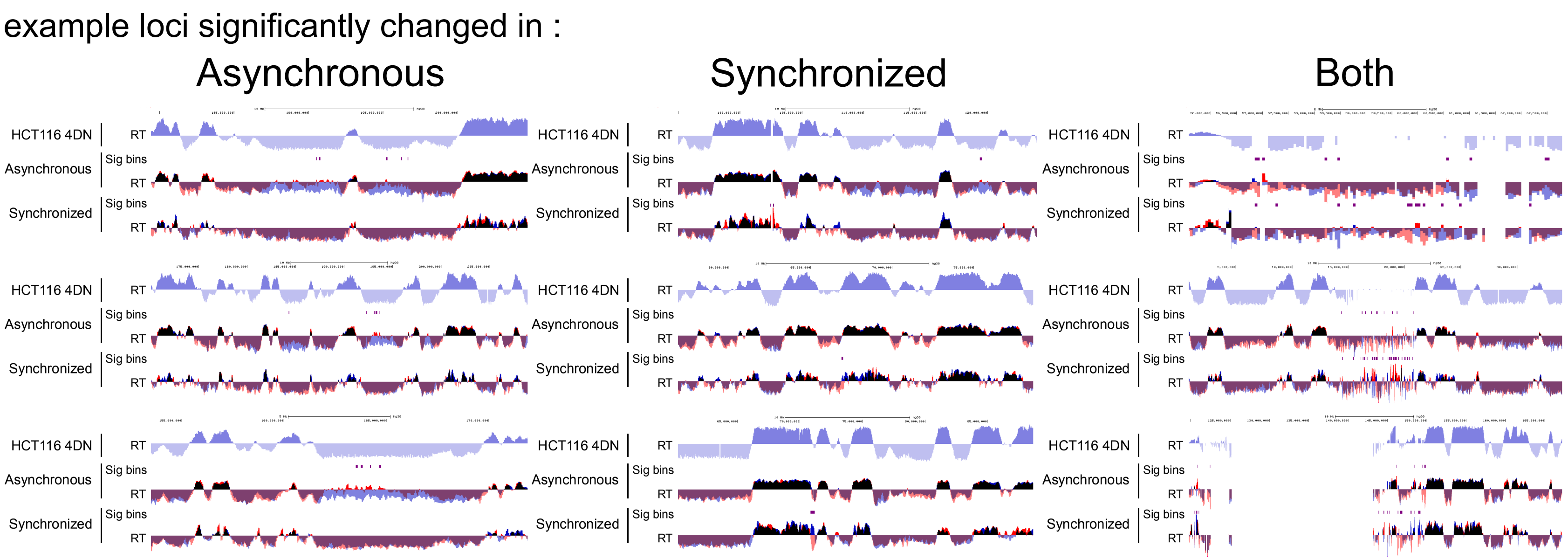


**Supplemental Figure 6. Loci with significant changes to replication timing, in asynchronous, synchronized, or both experiments.** Significance determined by FDR adjusted $p$ value < 0.05.

**Supplemental Table 1**. Source for genomic features annotated in HCT116 cells.

| Feature | Source |
|---|---|
| Super enhancers | (Hnisz *et al*, 2013) [45] |
| H3K9me3 | ENCSR179BUC |
| H3K9me2 | ENCSR555LYM |
| CTCF | ENCSR000BSE |
| H3K27Ac | ENCSR000EUT |
| H3K9Ac | ENCSR093SHE |
| H3K4me3 | ENCSR333OPW |
| H3K4me2 | ENCSR794ULT |
| H3K4me1 | ENCSR161MXP |
| H4K20me1 | ENCSR474DOV |
| H2AFZ | ENCSR227XNT |
| EZH2 | ENCSR046HGP |
| Dnase Hotspot | wgEncodeRegDnaseUwHCT116Peak (Sabo *et al*, 2006) [46] via UCSC Table Browser |
| CpG Island | cpgIslandExt via UCSC Table Browser |
